# Multimodal mapping of neural activity and cerebral blood flow reveals long-lasting neurovascular dissociations after small-scale strokes

**DOI:** 10.1101/2020.03.04.977322

**Authors:** Fei He, Colin Sullender, Hanlin Zhu, Michael R. Williamson, Xue Li, Zhengtuo Zhao, Theresa A. Jones, Chong Xie, Andrew K. Dunn, Lan Luan

## Abstract

Neurovascular coupling, the close spatial and temporal relationship between neural activity and hemodynamics, is disrupted in pathological brain states. To understand the altered neurovascular relationship in brain disorders, longitudinal, simultaneous mapping of neural activity and hemodynamics is critical yet challenging to achieve. Here, we employ a multimodal neural platform in a mouse model of stroke and realize long-term, spatially-resolved tracking of intracortical neural activity and cerebral blood flow in the same brain regions. We observe a pronounced neurovascular dissociation that occurs immediately after small-scale strokes, becomes the most severe a few days after, lasts into chronic periods, and varies with the level of ischemia. Neuronal deficits extend spatiotemporally whereas restoration of cerebral blood flow occurs sooner and reaches a higher relative value. Our findings reveal the neurovascular impact of mini-strokes and inform the limitation of neuroimaging techniques that infer neural activity from hemodynamic responses.

## Introduction

In healthy brains, local neuronal activity is strongly correlated, both spatially and temporally, with subsequent changes in cerebral blood flow (CBF). This close spatial and temporal relationship between neural activity and hemodynamics(*1, 2*), known as neurovascular coupling, is believed to be impaired in a wide spectrum of neurological and cerebrovascular diseases including Alzheimer’s disease, hypertension, and stroke(*3*). However, the longitudinal alteration of neurovascular coupling during disease progression and recovery remains largely unstudied. In this study, we use a multimodal, chronic neural interface in a mouse model of stroke to simultaneously record neural and hemodynamic activities and examine whether, when, and to what extent CBF correlates with the underlying neural electrical activity.

In ischemic stroke, the reduction of cerebral blood flow induces a complex series of pathophysiological events that evolve in time and space(*4*). The classical view is that local tissue perfusion dictates the neuronal response. Studies of middle cerebral artery occlusion (MCAO) in humans and animal models have established the ischemic threshold of CBF for irreversible neural damage at acute periods(*5, 6*). *In vivo* two-photon (2P) imaging in animal models has shown that suppression of CBF following MCAO induces structural impairment on axons(*7*), dendrites(*8*), and spines(*9*), and may be reversible depending on local reperfusion(*10*) shortly after the occlusion. Simultaneous measurements of CBF and evoked potentials in a rat model revealed variations of neurovascular coupling with the level of global cerebral ischemia(*11*). However, most of the studies were performed at acute phases, largely due to the challenges of repeatedly quantifying multiple neuronal and hemodynamic activities simultaneously with sufficient spatiotemporal resolutions over chronic periods. It remains to be understood how the interaction between CBF and neuronal activity unfolds over weeks and longer, and how it is spatially graded and temporally staged by the injury.

In this study, we spatially resolved neuronal electrical activity and CBF in the same brain regions and tracked their changes from the pre-stroke baseline to eight weeks after stroke. These measurements were enabled by a chronic multimodal neural interface that combined targeted photothrombosis, laser speckle contrast imaging (LSCI) of CBF, and intracortical neural recording of local field potential (LFP) and spiking activities using ultraflexible electrode arrays (Figure 1). We took advantage of our recently developed ultraflexible nanoelectronic threads (NETs)(*12*), which facilitate chronic optical imaging and afford long-lasting, stable recording of unit activities at minimal perturbation to the baseline neurophysiology(*12, 13*). We combined multi-shank, multi-depth neural recording with multi-exposure speckle imaging (MESI), a refinement of speckle contrast imaging that enables quantification of CBF for longitudinal and cross-animal comparisons with high resolution at large field of view (*14, 15*). By patterned illumination using a digital micromirror device (DMD), we induced targeted photothrombotic occlusion within individual or multiple surface arteriole branches to provide fine control over lesion location, size and onset time(*16*). In addition, by confining photodamage to arteries on the cortical surface, this modified technique produces a more graded penumbra than traditional photothrombosis(*17*). Finally, we performed all measurements including the induction of stroke on awake, head-fixed animals to remove the confounding impacts of anesthetic agents on neural activity, hemodynamics, and neurovascular coupling(*18–20*). Taking advantage of these technical improvements on multiple fronts, we investigated the following questions: how does neuronal activity change in response to CBF variations at different stages after small-scale strokes? Whether, when, and to what extent does the change of CBF dictate or reflect neural activity after the injury? These questions are important to understand the neurovascular impact of small-scale strokes and microinfarcts, which are thought to have a strong link to vascular dementia(*21, 22*). Furthermore, the deficits and restoration of CBF in the peri-infarct tissue are often used as surrogate measures to evaluate neuronal deficits and to predict recovery at the acute and subacute phases(*23*). Should the correlation between neural and CBF activities alter with the level of ischemia or with time and space, we must consider the details of the alteration for the application and precise interpretation of clinal neuroimaging techniques such as functional magnetic resonance imaging (fMRI) (*24, 25*) that use hemodynamic responses to infer neural activity.

**Fig. 1.**
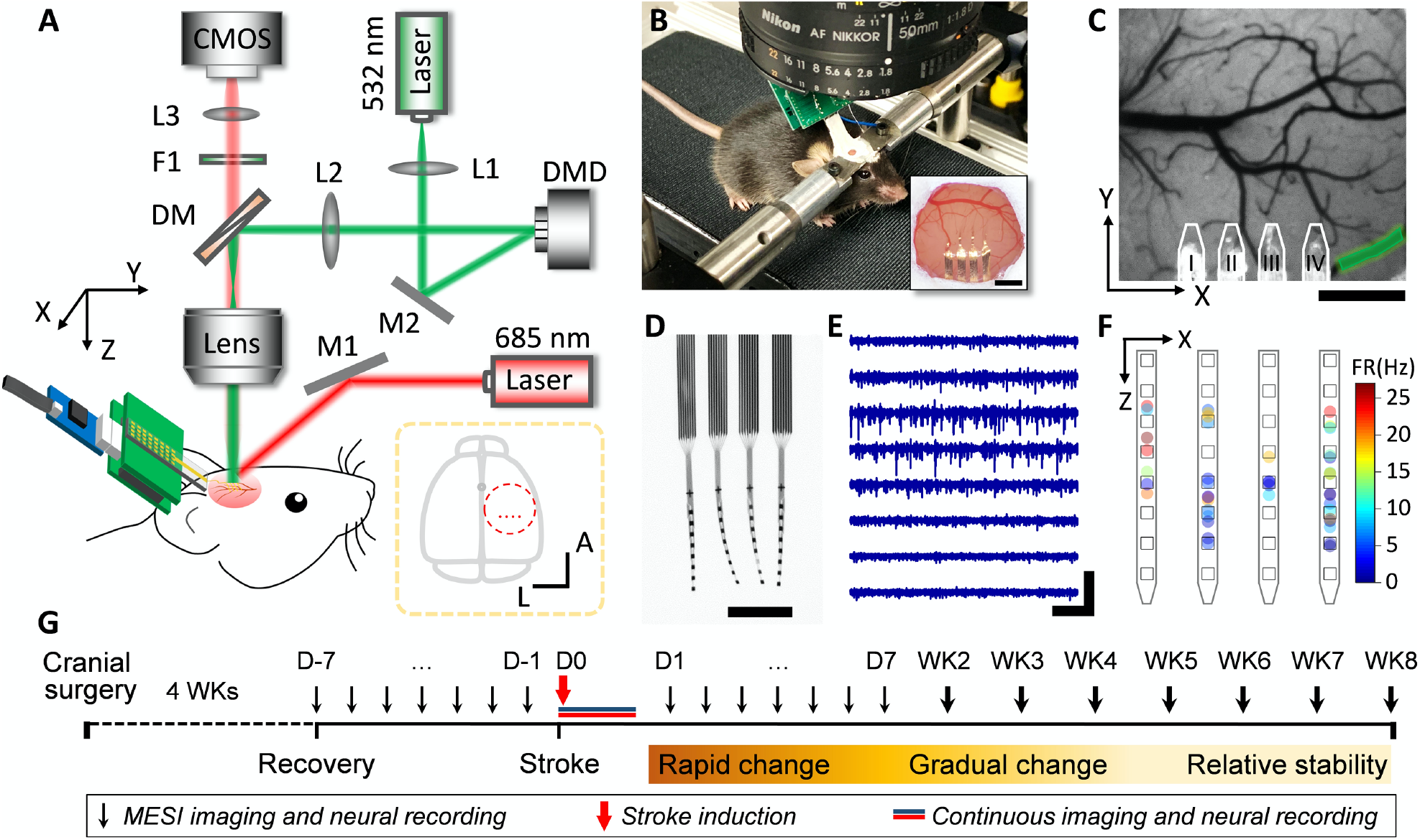
The multimodal neural interface for simultaneous mapping of CBF and neuronal electrical activity. **(A)** Schematics of the multimodal measurement. The optical paths for targeted photothrombosis (532 nm, green) and the speckle imaging (685 and 785 nm, red) were sketched. DM: dichroic mirror, F1: filter, L1-3: lenses, M1-2: mirrors. Inset: dashed circle shows the anatomical location of the cranial window relative to bregma (gray dot). Red dots indicate the implantation sites of the 4-shank probes. A: anterior; L: lateral. **(B)** Photograph of a mouse on a treadmill under the imaging apparatus implanted with NETs and a cranial window. Inset: photo of the cranial window at 10 weeks after surgery. Photo credit: Fei He, Rice University. **(C)** A representative LSCI of pre-stroke CBF, where the green shade highlights the arterioles for targeted photothrombosis, and the white lines outline the NETs. **(D)** Photograph of a NET array (4 × 8 recording sites) in water showing its ultraflexibility. Photo credit: Fei He, Rice University. **(E)** Typical recording traces (300 Hz high-pass filtered) from 8 recording sites on one NET shank. **(F)** Spatially resolved spike activities from a 5-min representative recording session using a 4 × 8 NET array. Squares sketch individual contacts; dots are color-coded to present spike rates, whose locations are estimated as the averaged locations of the contacts weighted by the spike magnitudes they detect. **(G)** Experimental procedure and timeline. Scale bars: 500 μm (**B** (inset), **C** and **D**); 400 μV (vertical in **E**) and 200 ms (horizontal in **E**).

## Results

All experiments involved awake adult C57/BL6J mice that were acclimated to head fixation (**Materials and Methods**). We employed a modified cranial window technique and implanted multi-shank NET arrays, which permitted spatially resolving neural activity spanning the depth and lateral locations under the optical window (Figure 1A–1F, representative recording from 4-shank, 32-contacts shown in **Figure S1**)(*12, 13*). The experiments started at least four weeks after the cranial surgery, which allowed for the recovery of nearby vasculature from the implantation damage (*12, 13*). We induced targeted photothrombosis and occluded one or a few branches of descending arterioles to induce small-scale, focal ischemia in the sensory cortex of awake mice. We repeatedly performed MESI of CBF and electrophysiological recordings in the same brain regions. Because the peri-infarct tissue is expected to go through rapid neural and vascular plasticity after stroke, it is more challenging to repeatedly record from the same individual neurons than in healthy brains. Therefore, we performed population-based quantification for neural recording including LFP spectral weight as well as single- and multi-unit spiking rate without chronically tracking individual neurons’ activities. The measurements included multiple pre-stroke baseline sessions, the acute stroke session, and the chronic periods for up to 8 weeks after stroke. The experimental time course was determined by our pilot studies showing that stroke-induced neurovascular changes had subsided and all variables we longitudinally tracked were relatively stable by week 6. One animal was terminated at week 5 due to skull regrowth under the cranial window that induced clouding and impeded quantification of CBF from speckle imaging. The exclusion of animals and locations are detailed in **Table S1**. We tailored the measurement frequency to the anticipated changes in neuronal and CBF responses. We sampled more frequently immediately after stroke for the most dynamic phase and relaxed the sampling rate in the chronic periods when relative stability was expected (Figure 1G).

### Neurovascular coupling weakens with ischemia at acute periods after stroke

We simultaneously imaged CBF and recorded neural activity in awake animals shortly before, during, and after green light illumination for targeted photothrombosis. This allowed us to monitor the infarct formation *in vivo* (**Figure S2**) and to capture the variations of CBF and neuronal electrical activity from pre-stroke baseline to during the formation of the lesion and after. We chose to selectively occlude arteriole branches in the imaging field of view (Figure 2A) and harnessed the spatially graded ischemia induced by this refined method of artery-targeted photothrombosis(*17*). By implanting 2D NET arrays at multiple distances from the infarct, we recorded at multiple locations and sampled varying levels of ischemia (Figure 2B–2D). We determined the single- and multi-unit firing rate and LFP power density at multiple frequency bands recorded at individual NET contacts. We quantified the relative values of parenchymal CBF from LSCI (**Materials and Methods**) in the tissue where NETs were implanted. Representative time traces from 90-min continuous recording and imaging at one location is shown in Figure 2E, 2F and the full-field imaging and recording is shown in **Movie S1**. Notably, at the normal brain state pre-stroke and under mild ischemia, the variations in neural activity were tightly coupled to the variations in CBF. However, after parenchymal CBF significantly reduced following the spontaneous occurrence of peri-infarct depolarizations (PIDs), variations in neural activity led to little variations in CBF. Correspondingly, the scatter plots from multiple animals (N = 10) and brain locations (n = 39) show two distinct clusters: positive linear correlation between relative CBF (rCBF) and relative LFP (rLFP), as well as between rCBF and relative spike firing rate (rFR) for no and mild ischemia, and no correlation after CBF had dropped significantly (Figure 2G, 2H and **Figure S3A-S3D**). Importantly, although some recording sites detected prolonged electrical silencing after PIDs, most sites still recorded substantial neural activities after the membrane potential recovered. However, their correlation with CBF changes was lost under ischemia. Increases in neural activity no longer consistently led to increases of CBF, and spontaneous restoration in CBF was disassociated with the recovery of neural activity.

**Fig. 2.**
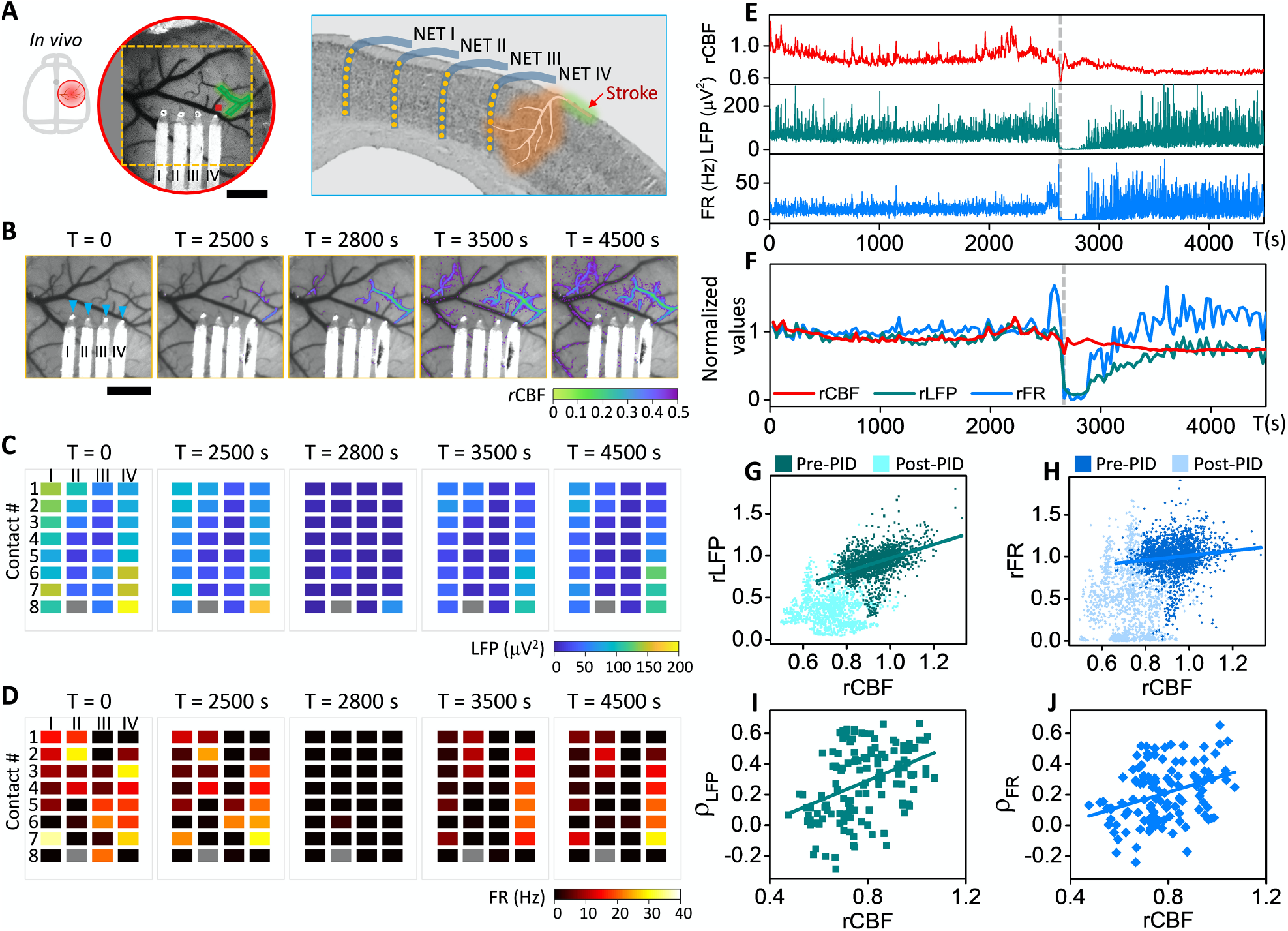
Neurovascular coupling weakens with ischemia in acute periods after stroke. **(A)** Sketch of the skull (left), LCSI of CBF (middle) and sketch on a coronal section (right) showing the locations of the cranial window, 4 NET shanks and 32 contacts in **(B–D)**, and the targeted arteriole for photothrombosis (shaded in green). Red square marks the ROI used to quantify rCBF in **(E, F)**. **(B)** Time series of LSCI of CBF at the acute stroke session. Overlaid color map shows the area of rCBF < 50% baselined against pre-photothrombotic images. Blue triangles mark the NET implantation sites. **(C, D)** Time series of LFP at 60–110 Hz and spike firing rate (FR) from all NET contacts at the acute stroke session. Time points match those of **(B)**. Roman and Arabic numbers mark the NET shanks and contacts along the depth, respectively. One disconnected contact (II-8) is plotted in grey. **(E, F)** Simultaneous measurement of LFP and FR recorded by contact IV-4 and CBF from the ROI in **(A)** during and following photothrombosis. Normalized and 30s-averaged values were used in **(F)** to visualize the neurovascular dissociation. Dashed lines mark a PID event that was excluded from the following analysis. **(G, H)** Scatter diagrams of rCBF and rLFP at 60-110 Hz **(G)**, and rCBF and rFR **(H)** for all locations (n = 39) and animals (N = 10) showing the positive linear correlation (darker color) breaks down as ischemia worsened (lighter color) after a PID occurred. **(I, J)** Correlation coefficients computed between rCBF and rLFP **(I)**, and between rCBF and rFR **(J)**. Pearson’s correlation was applied: ρ = 0.42, p-value = 4e-94 **(G)**; ρ = 0.12, p-value = 8e-9 **(H)**; ρ = 0.40, p-value = 3e-6 **(I)**; ρ = 0.35, p-value = 1e-4 **(J)**. Scale bars: 500 μm (**A** and **B**).

To quantify the alteration of neurovascular coupling due to ischemia, we computed the correlation coefficient between neural activity and parenchymal CBF for sub-sessions of 1000 s in duration (**Materials and Methods**). To determine the level of ischemia in the sub-session, we averaged the parenchyma CBF over time and normalized it to the pre-stroke baseline value. Sub-sessions in which any PIDs occurred were excluded from the analysis because PIDs lead to neuronal electrical silence and waves of hypoperfusion(*26*). Figure 2I, 2J and **Figure S3E, S3F** show the scatter diagram of the correlation coefficient between CBF and neural electrical activity computed at different frequency bands. A linear correlation was found between CBF and neural-CBF correlation coefficient for all frequency bands of LFP and spike activity (ρ = 0.35 for spike rate; ρ = 0.37 for multi-unit band at 0.3k–3k Hz; ρ = 0.44 and ρ = 0.34 for LFP at 60–110 Hz and 30–60 Hz. p-value < 0.001 for all datasets). This suggests that neurovascular coupling weakens with the severity of ischemia. When CBF declined below 40% of the pre-stroke baseline, the neurovascular coupling coefficient reduced to zero. Because LSCI provides depth-integrated CBF, we used depth-averaged neural data to best match the depth profile of CBF. We examined a variety of depth profiles of neural activity, including using neural activity recorded by the single, shallowest contact, by the contact that measured the largest correlation coefficient at pre-stroke, using the averaged value recorded from a subset of contacts, and using all contacts on the same shank (**Figure S4**). We confirmed that the coupling coefficients between neural activity and CBF showed consistent dependence on the level of ischemia irrespective of the choice of the depth profile.

### Neurovascular disassociation is long-lasting and most severe at sub-acute periods

We longitudinally tracked the spontaneous restoration of CBF and recovery of neural activity after stroke (Figure 3 and **Movie S2**). We detected a significant discrepancy between them that became the most severe in the sub-acute phases (up to two weeks after stroke) and lasted into the chronic periods. First, the restoration of CBF and recovery of neural activity have distinct time courses. Longitudinal MESI of CBF (baselined against pre-stroke measurements) showed that blood flow in the peri-infarct cortex decreased significantly after stroke and improved shortly after. Reperfusion in the occluded vessels and nearby tissue occurred promptly in all animals at a time course of 1–3 days after targeted photothrombosis (Figure 3C). This was, however, accompanied by significantly suppressed neural activity measured by the reduction in both LFP and FR from simultaneous electrical recording in the same brain region (Figure 3D-3F). Neural activity remained inactive for a more extended period (*e.g.* Day 7 in Figure 3D-3F) even after blood flow was fully restored (Figure 3C), and gradually improved over time. Secondly, the variations in magnitude were different for CBF and neural activity. While hyperperfusion was pronounced in all animals, neuronal hyperexcitability(*27, 28*), which manifested as higher-than-baseline spike rate and upshift in the frequency distribution of spiking activities, was completely absent (Figure 3D-3F) in 7 out 10 animals. In the other 3 animals where hyperexcitability was detected, it was spatially sparse and detected only by 13 contacts in 8 shanks out of 90 contacts in total. As shown in **Figure S5**, hyperexcitability occurred in microscopic regions interspersed among regions of hypo-excitability and relatively stable firing patterns. In addition, it occurred at a delayed onset time of a few days to 2 – 3 weeks later from the occurrence of hyperperfusion. Finally, after the dynamic changes subsided with time, neural activities often remained substantially reduced even when CBF restored to a value close to the pre-stroke baseline (Day 28–55 in Figure 3C-3F).

**Fig. 3.**
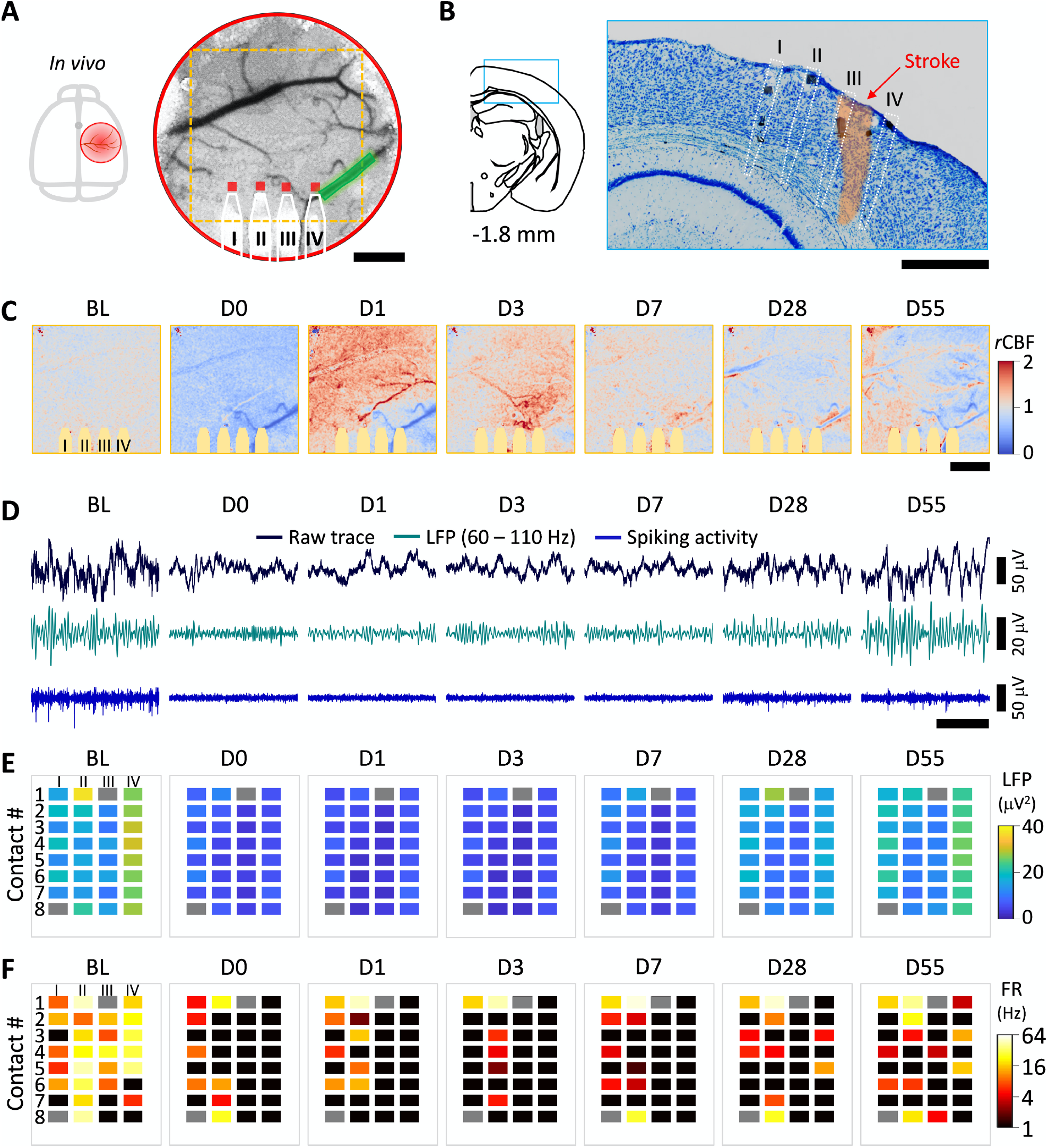
Neurovascular disassociation lasts chronically after stroke. **(A)** Sketch of the skull and LSCI of CBF before stroke showing the location of the cranial window, the implanted NET shanks (highlighted by white solid lines), the targeted arteriole for photothrombosis (green shading), and the ROIs (red squares) used in Fig. 4A. Dashed squares show the area in **(C)**. **(B)** Nissl-stained coronal sections showing the NET implantation locations (white dashed lines) and the infarct area (orange shading). **(C)** MESI of CBF at multiple temporal coordinates before and after stroke. Colormap is baselined against the averaged pre-photothrombotic values to highlight both hyperperfusion (red) and hypoperfusion (blue) regions. The yellow shades mark the regions under NETs, which were excluded from the quantification of CBF. Days after stroke are marked on top. BL: pre-stroke baseline. (**D**) Representative, single-channel recordings from contact IV-5 at the same temporal coordinates as in (**C**). Black: raw trace; green: LFP at 60-110 Hz; blue: spiking activity. **(E, F)** LFP at 60**–**110 Hz (**E**) and spike rate **(F)** using all NET contacts. Two disconnected contacts are plotted in grey. Restoration of CBF occurred on Day 1, while neural activity remained significantly suppressed several days after. Scale bars: 500 μm **(A–C)**; 200 ms (horizontal in **D**).

We determined the relative values of all measurements normalized to the averaged pre-stroke baselines and quantified the longitudinal variations in CBF, LFP and FR at each region of interest (Figure 4A). Their relationship strongly depended on the temporal stage. At the pre-stroke phase (7 consecutive days), the inter-day variations in both LFP and CBF were small, and their values remained close to the baseline. At the sub-acute phase (Day 1–14 poststroke) when hemodynamic and neural activities went through the most dynamic changes, the variations in LFP and CBF were both significant, but their variations were decoupled. At the chronic phase when the dynamic responses subsided (week 3–8 after stroke), neural activity progressively recovered while CBF remained relatively stable with day-to-day variations on par with that detected at the pre-stroke phase. The CBF-LFP scatter diagram from all animals (N = 10) and locations (n = 37) had a narrow distribution symmetrically centered at the averaged baseline values at the pre-stroke phase, became widely distributed at the subacute phase, and re-clustered at the chronic phase but nevertheless had a much wider distribution than at the pre-stroke phase (Figure 4B). The CBF-FR scatter diagrams showed similar changes at the three phases, but were more distributed at the sub-acute and chronic phases (Figure 4C). Consistently, the distribution of each parameters was the widest at the sub-acute phase (Figure 4D–4F). Notably, there was no statistically significant difference between the values of CBF at baseline and at the sub-acute phase, both of which were larger than the CBF at the chronic phase. However, LFP and FR at the sub-acute phases were substantially suppressed. Correspondingly, the averaged value of r(CBF-LFP) and r(CBF-FR) was zero at baseline, and shifted to positive values at the subacute and chronic phases, suggesting that CBF restored to higher relative values than neural activity during spontaneous recovery (Figure 4G, 4H). r(CBF-LFP) and r(CBF-FR) were the largest at the subacute phases, suggesting that the neurovascular disassociation was the most significant at this stage. LFPs at other frequency bands showed qualitatively consistent results in their distributions and relationships with CBF at the three temporal stages after stroke (**Figure S6**).

**Fig. 4.**
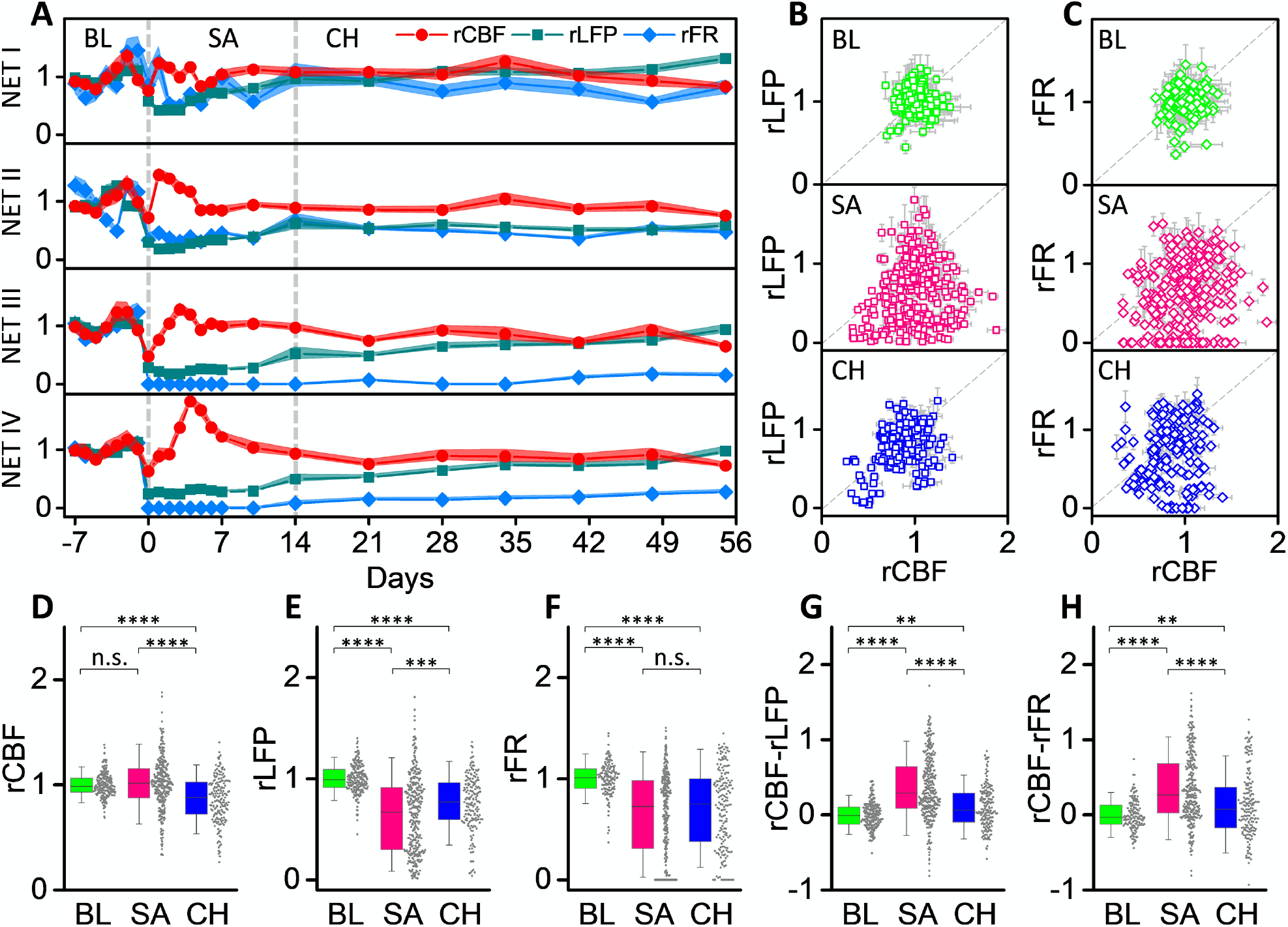
Neurovascular disassociation is the most severe at the sub-acute phase. **(A)** Relative CBF, LFP and spike rate at the 4 NET locations in Fig. 3 for 9 weeks baselined against pre-stroke values. Solid dots present the mean values and shading shows the STD. Neurovascular disassociation is the most evident at the subacute phase, where the difference between rCBF and rLFP/rFR is the largest. **(B, C)** Scatter diagrams between rLFP and rCBF **(B)**, and between rLFP and rFR **(C)** in three phases: baseline (BL), consecutive 7-days daily measurements before stroke; subacute (SA), Day 0–14 post stroke; and chronic (CH), week 3–8 post-stroke (location n = 37 and animal N = 10). **(D – H)** Box plots showing the relative values of CBF **(D)**, LFP **(E)**, spike rate **(F)**, and their difference CBF-LFP **(G)** and CBF-spike rate **(H)** at the three phases: baseline (green); sub-acute (magenta); chronic (blue). All data plotted as dots. Significant levels shown as: n.s., no significance; **, p-value < 0.01; ***, p-value < 0.001; ****, p-value < 0.0001.

### More severe ischemia leads to longer-lasting neural deficits and neurovascular dissociation that are best predicted by acute neural activity

We varied the degree of ischemia by controlling the number and dimension of the arteriole branches for targeted photothrombosis. We measured the spatial extent of the tissue under ischemia and the infarct size in two means: the area under 20% CBF of the pre-stroke baseline from *in vivo* MESI of CBF at the end of the acute session (Figure 5A), and the lesion volume identified in postmortem Nissl staining at 8 weeks after photothrombosis (Figure 5B). A previous study showed that tissue damage identified from histological analysis paralleled regions of severe CBF deficits and that MESI estimates of ≤20% baseline CBF served as a reliable *in vivo* estimate of the ischemic core (*17*). We used the area under severe CBF deficits in MESI as a measure of infarct dimension and examined its correlation with the severity of reduction in neurovascular coupling. To capture both the magnitude and duration of the neurovascular changes induced by ischemia, we integrated with time the relative deficit in CBF and LFP baselined against pre-stroke values: ***D***_***CBF***_ = ∫(**1** − ***rCBF***)***dt*** and ***D***_***LFP***_ = ∫(**1** − ***rLFP***)***dt***. The difference of the two values, ***D***_***CBF***_ − ***D***_***LFP***_, characterizes the duration and magnitude of neurovascular disassociation. We did not detect a statistically significant correlation between ***D***_***CBF***_ − ***D***_***LFP***_ and area of severe CBF deficit (**Figure S7A**, ρ = 0.21; p-value = 0.25). We attribute this lack of correlation to the fact that the measurements were performed in the peri-infarct tissue at varying distances from the infarct core. Consequently, each measurement sampled different levels of ischemia due to the spatial variation of ischemia within animals and lacked the consistency on ischemic severity for cross-animal comparisons.

**Fig. 5.**
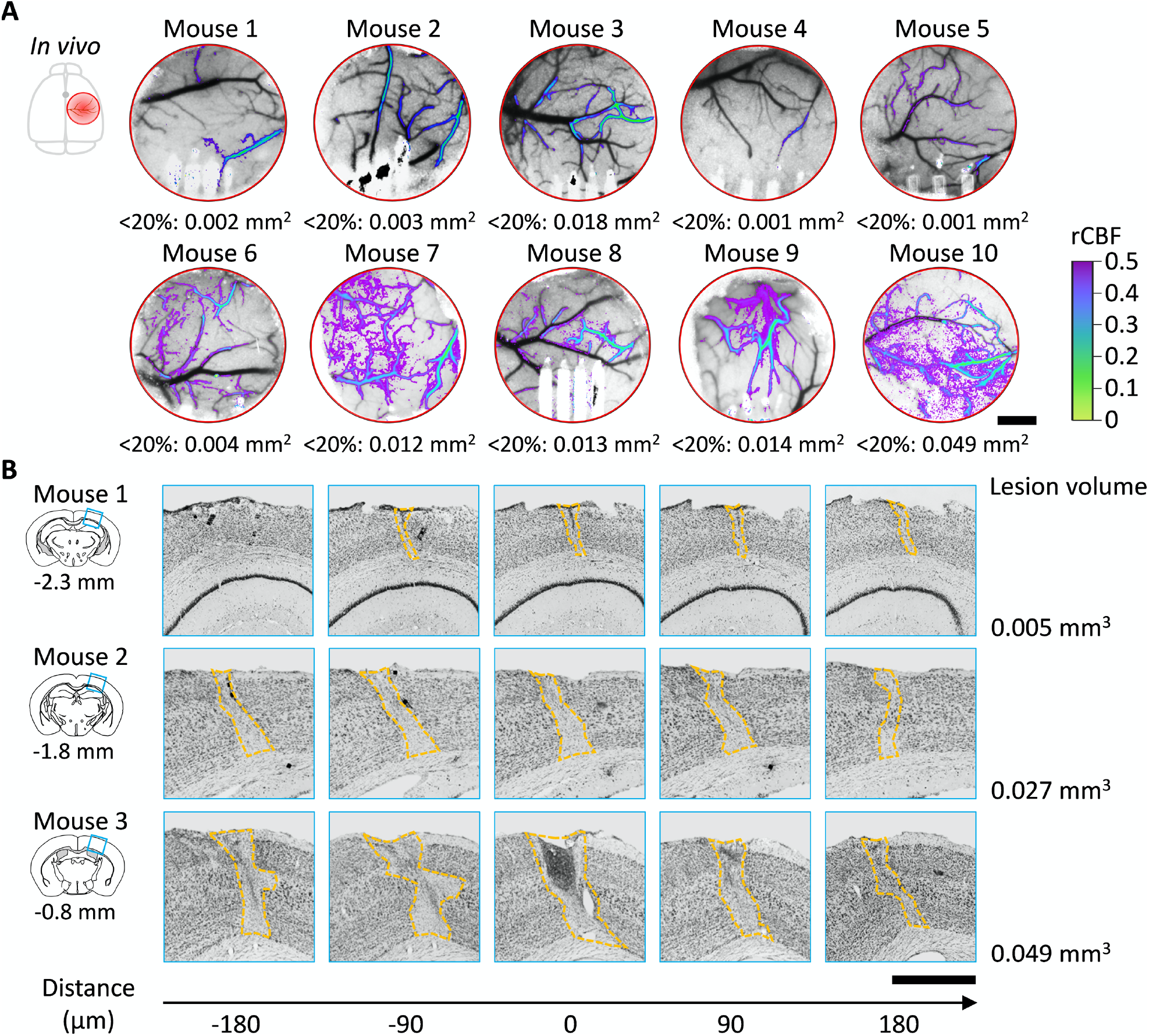
*In vivo* and histological measurements of infarct dimensions. **(A)** LCSI of CBF at the end of the acute session showing the spatial extent and severity of ischemia in all animals. Colormap: overlay of rCBF < 50% baselined against pre-photothrombotic images. Area under rCBF < 20% is denoted under each image. **(B)** Nissl staining of the infarct tissue at 8 weeks after stroke for three animals as shown in **(A)**. The coronal section templates indicate the infarct positions posterior to bregma. Numbers on the bottom are the relative distance to the templates. Scale bars: 500 μm (**A** and **B**).

To account for the spatial gradient within individual animals and the varying scales of ischemia across animals, we employed two location-specific predictors as measures of the initial injury: CBF_0_ and LFP_0_ measured at the locations where NETs were implanted on Day 0 immediately after the lesion was fully formed. We first examined the final neural and hemodynamic outcomes measured as LFP_end_ and CBF_end_ at 6 weeks poststroke at the same locations (animal number N = 9; location n = 33). LFP_end_ is linearly correlated with both CBF_0_ (ρ = 0.41; p-value = 0.02, Figure 6A) and LFP_0_ (Pearson’s correlation: ρ = 0.64; p-value = 4.1e-5, Figure 6B), but the correlation with LFP_0_ is stronger than with CBF_0_. In contrast, CBF_end_ has no correlation with either acute CBF_0_ (ρ = 0.20; p-value = 0.25, **Figure S7B**) or LFP_0_ (ρ = 0.15; p-value = 0.41, **Figure S7C**). This is consistent with the observation that CBF restored to pre-stroke baselines for a wide range of strokes in this study, but LFP recovery was incomplete after more severe ischemia.

**Fig. 6.**
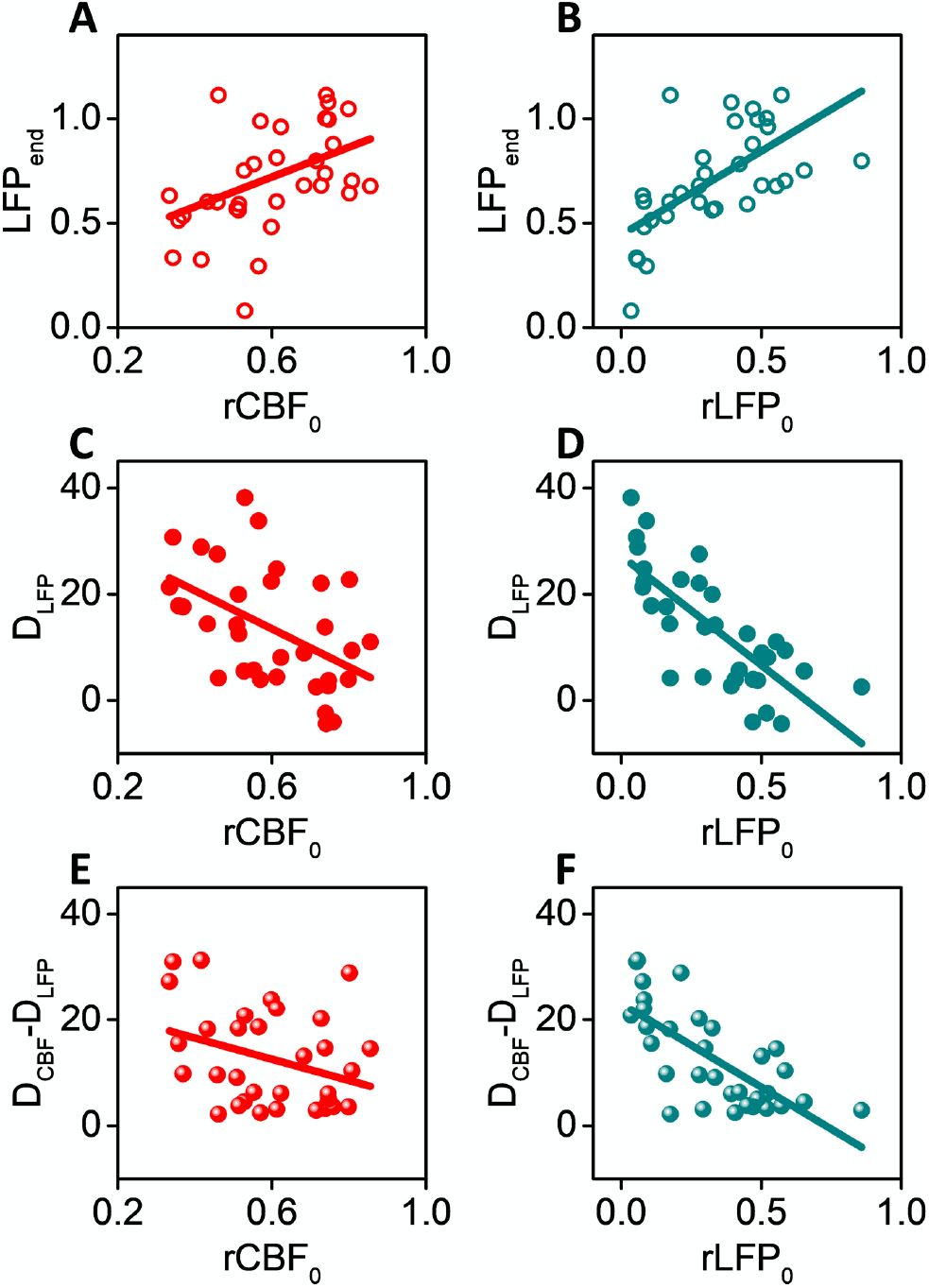
More severe ischemia leads to longer-lasting neural deficits and neurovascular disassociation. **(A-F)** Pearson’s correlation using acute values of CBF and LFP as predictors for LFP at week 6 post stroke (ρ = 0.41, p-value = 0.02 in **(A)**; ρ = 0.64, p-value = 4.1e-5 in **(B)**), for time-integrated neural deficits (ρ = −0.49, p-value = 0.004 in **(C)**; ρ = −0.76, p-value = 2.3e-7 in **(D)**), and for time-integrated neurovascular disassociation (ρ = −0.33, p-value = 0.06 in **(E)**; ρ = −0.71, p-value = 3.3e-6 in **(F)**). The Pearson’s correlation coefficient is larger and p-value is smaller when using LFP_0_ as the predictor than using CBF_0_ for all three variables.

We further examined the duration and magnitude of neurovascular disassociation by using the time-integrated values ***D***_***CBF***_, ***D***_***LFP***_, and ***D***_***CBF***_ − ***D***_***LFP***_ (Figure 6C – 6F). Markedly, although global measure of CBF deficits (Figure 4) had little correlation with local-specific neural or hemodynamic outcomes, local values of CBF_0_ was weakly correlated with ***D***_***LFP***_ (ρ = −0.49; p-value = 0.004) and ***D***_***CBF***_ − ***D***_***LFP***_ (ρ = −0.33; p-value = 0.06). Furthermore, LFP_0_ demonstrated stronger correlation with both ***D***_***LFP***_ (ρ = −0.76; p-value = 2.3e-7), and ***D***_***CBF***_ − ***D***_***LFP***_ (ρ = −0.71; p-value = 3.3e-6). These results suggest that more severe ischemia leads to longer-lasting, higher magnitude neurovascular deficits and disassociation. However, because the strong correlation between neural activity and CBF breaks down due to ischemia, acute neuronal deficits such as the reduction in LFP rather than acute reduction of CBF better predict the long-term neuronal outcome for the small scales of strokes in this study.

## Discussion

Extensive studies(*7–10, 27–29*) using state-of-the-art imaging and electrophysiological tools have increased our knowledge on the pathological neurovascular responses to ischemic injury, which is of great importance for both clinical and basic neuroscience. However, comparable approaches to combine optical imaging and electrophysiology in the same brain region and to simultaneously track neural and hemodynamic changes into chronic time scales are currently minimal. In this work, we take advantage of a multimodal, chronic neural platform that combines speckle imaging of cerebral blood flow and spatially resolved intracortical neural recording to unveil the neurovascular impact of varying scales of focal ischemic injuries. We identify pronounced dissociations between hemodynamic and neural responses that are injury dependent and time sensitive. Our results provide meaningful complementation to previous studies to determine multiple neurophysiological parameters simultaneously in the extended time courses and at various sizes of injuries that bridge small-scale strokes and individual cortical microinfarcts.

In acute sessions immediately after infarcts are induced, the neurovascular response has mostly been studied in anesthetized animals using sensory evoked responses(*11, 30*). We measured spontaneous neural activity in the resting state of awake animals and detected a similar threshold for electrical silence at CBF values of 40% of the pre-stroke baseline. We fill the knowledge gap in prior studies by revealing the progressive reduction of the neurovascular coupling by ischemia at levels above this threshold. Notably, moderate levels of ischemia do not causally attenuate neural activity. Instead, spontaneous variations in neural activity become less correlated with subsequent changes in CBF.

Neuronal activity after stroke during chronic time courses that are commensurate to this study has been carefully investigated using *in vivo* imaging. Voltage-sensitive dye imaging in adult mice reveals that somatosensory maps lost to photothrombosis are replaced over several weeks by new structural and functional circuits(*31*). During this period of recovery, dendrites and spines are initially lost, but recover in a distance-dependent manner from the infarct(*31, 32*) with spine turnover surpassing the baseline in more proximal regions. Simultaneous measurements of spine dynamics and blood flow from surrounding capillaries before and after unilateral MCAO suggest that local hemodynamics dictate long-term dendritic plasticity in the peri-infarct cortex(*33*). Our study agrees with these studies on the time scale of several weeks and longer for the recovery of neuronal activity, and the dependence of the final neuronal outcome on the initial ischemic severity. But there are also important differences between those studies and the present one. First, we report significant, long-lasting disassociation between neuronal electrical activity and local parenchyma blood flow that was not reported before. Particularly, reperfusion does not lead to the recovery of neural activity immediately. There is a prolonged delay in time for neural electrical activity to approach the normal levels after CBF has restored or surpassed the pre-stroke baselines. This is in stark contrast to the acute imaging studies where rapid reversible changes in dendritic spine structure were gated by reperfusion(*9*). Secondly, although local CBF immediately after stroke is weakly correlated with final neuronal recovery, the deficits in LFP at the acute phase is a better predictor of final neuronal outcome and the recovery process. Finally, previous studies reported enhanced neuronal connectivity and spine density surpassing baseline values that depended on the distance from the infarct (*31, 32*). Our study detects long-lasting neural deficits with sparse neuronal hyperexcitability that does not present distance or depth dependence.

Multiple factors may contribute to these differences in findings. First of all, we longitudinally tracked both CBF and neuronal electrical changes after initial injury at identical brain regions and multiple temporal coordinates. This allows us to unveil the neurovascular dissociations that are the most pronounced at the subacute phases after stroke. This crucial period of time was not put under the spotlight in previous studies that mostly focused on the initial states and final states for concurrent measurements of CBF and neural activity(*31, 32*). Secondly, differences in results among previous studies and ours could be attributed to differences in the size of the penumbra created with the stroke models. We use a refined photothrombotic stroke model that selectively targets a single or a few branches of penetrating arterioles without non-discriminant tissue damage inclusive of microvasculature and veins. This created a much smaller, focal lesion than MCAO(*34*). Furthermore, this refined model of photothrombosis produces a more graded vascular penumbra than traditional photothrombosis while maintaining the ability to create localized infarcts(*17*). Thirdly, we sample a different portion of the cortex both in depth and in distance from the infarct compared with 2P imaging studies. The NET recording arrays are typically implanted 100 μm – 1 mm from the lesion and span a depth of 600 μm. 2P imaging experiments, in comparison, are mostly performed on the superficial layer of the cortex and at least a few hundreds of microns away from the infarct due to compromises on optical clarity from tissue injury. Fourth, we directly recorded spontaneous neuronal electrical activity including LFP, single-unit action potentials, and multi-unit spikes in awake brains. This may be indirectly related to the structural plasticity manifested as dendritic spine turnover and density from a single layer of neurons, or functional connectivity induced by evoked stimulation and examined through post-mortem histology.

The conventional view of brain injury is that the magnitude of the deficit is coarsely correlated to the infarct size when controlled for other influences including age, morbidity and stroke locations(*35*). However, it is unclear whether it holds for microscopic ischemic injuries such as microinfarcts(*36, 37*) given the inherent nonlinearity of ischemic thresholds(*5, 6*), and how the deficits manifest in multiple neurophysiological measurements and evolve with time. Previous studies suggest that cortical microinfarcts follow a prolonged time course compared to larger regional infarcts(*21*), induce widespread deficits in a much larger volume of tissue than the microinfarct core(*38, 39*), and are insufficiently detected by MRI(*39*). We supplement these studies by varying the scales from ministrokes that span a few mm^2^ to individual cortical microinfarcts and tracking their neurovascular impacts longitudinally *in vivo*. We show that the time course and magnitude of the injury development depend on the initial injury, but the initial hemodynamic deficits and infarct sizes are not the best predictors for the tissue outcome and recovery duration. Prompt reperfusion to values approaching and exceeding pre-stroke baselines is accompanied by long-lasting neural deficits, which is consistent with the short detection window of microinfarcts in MRI and the long duration of functional impairments(*39*). The long duration of the neuronal recovery is consistent with the long-term dendritic plasticity reported for larger strokes at more distant sites(*29*), which suggests that the intervention time window is much longer than the subacute phases for the ischemic injuries of multiple scales in this study.

We did not measure motor deficits in this study. Recent studies showed that the artery targeted photothrombosis of similar and larger sizes than those induced in this study led to forelimb impairments in skilled reaching tasks but were difficult to detect using less sensitive behavioral assessments(*17, 40*). Further investigations that combine sensitive behavioral assessments and simultaneous measurements of neuronal structural changes and electrical activities will help to elucidate the underlying mechanisms of functional recovery.

Taken together, we demonstrate a multimodal neural platform capable of spatially resolving and longitudinal tracking multiple neurophysiological parameters in abnormal brain states. We reveal spatiotemporally varying neuronal and hemodynamic deficits after small-scale strokes that extend well beyond the lesion site and into chronic time scales. The long-lasting, injury-dependent neurovascular dissociation we unveil in this study indicates that hemodynamic parameters should be used with caution to infer neural activity for ischemic brain states. It raises the demand for direct neurological interrogations to evaluate brain impairment and recovery more accurately.

## Materials and Methods

### Subjects

Animal use and experimental design adhered to STAIR criteria and ethical animal welfare. 12 adult, male C57BL/6J wild type mice (4 – 6 months) were used in all experiments. They were housed one to four per cage at a 12-hr, 7:00 to 19:00, light-to-dark cycle. All animals received standardized cage supplementation (cage enclosures, nesting material, and objects to gnaw) with water/food *ad libitum*. All animals received one surgery during which a NET array was implanted intracortically and a chronic cranial window was mounted on the skull. 1 NET shank failed to be implanted intracortically, resulting a total of 47 locations in 12 animals. 2 animals (8 locations) were excluded from the acute-session data analysis due to abnormally high level of locomotion that induced motion-related artifacts and interfered with quantification of CBF and neural activity. 2 animals and 2 additional locations were excluded from the longitudinal studies in Fig. 4 due to post-stroke complications including animal loss and NET loss. One additional animal was excluded from Fig. 6 because of cranial window clouding at week 5 that resulted in earlier termination of the experiment. The experimental animal numbers are summarized in **Table S1**. Data were collected in a blinded manner whenever possible. In particular, the operator that induced photothrombosis and quantified their severity at acute phases was blinded to all following neural, hemodynamic and behavioral assessments. "Batch" effects were minimized by running staggered cohorts of equal numbers of animals per condition at a time with randomized condition assignment. All data analysis was performed using standardized procedures and largely automatic protocols with minimum human input(*41*). All experimental procedures were accordance with the National Institutes of Health Guide for the Care and Use of Laboratory Animals. All procedures have been approved by Institutional Animal Care and Use Committee at Rice University (protocol 1486756) and at the University of Texas at Austin (protocol AUP-2016–00257).

### Sample size estimation

Sample size was estimated using the pilot data from the first three animals in the study to detect an effect size equal to or smaller than 15% at a power of 0.8 in two-side *t* test and ANOVAs. The sample size was computed for all variables (CBF, LFP and spike rate) and the largest sample size needed (for the variable of spike rate, n = 33) was used in the study. The total animal number also included about 25% overage to counter attrition for surgery and post-stroke complications, which is detailed in **Table S1**. Power calculation was also performed after experiments to verify that the power was above 0.9 for all analysis that involved CBF and LFP, and above 0.8 for all analysis that involved spike rate.

### NET device fabrication and assembly

The NET devices were fabricated using planar photolithography fabrication methods similar to those previously reported(*12*). After a nickel metal release layer was deposited on a glass substrate (Soda Lime Glass 100-mm DSO 550-μm thick, University Wafer), SU-8 photoresist (SU-8 2000.5, MicroChem) was used to construct the bottom insulating layer. Interconnects were then patterned by depositing 120 nm gold. After a second insulating layer of SU-8, contacts with dimension of 30 μm × 30 μm were patterned by depositing 120 nm gold, electrically connected with the interconnects through “via”s on the top SU-8 layer. After device fabrication, a connector (36 Position Dual Row Male Nano-Miniature,. 025"/. 64 mm, Omnetics) was mounted on a home-build PCB adaptor board connected with the glass substrate. The flexible section of the devices was then released from the substrate by soaking in nickel etchant (TFB, Transene Company) for 4 – 8 hrs at 25 °C. Poly(3,4-ethylenedioxythiophene) (PEDOT) polymer coating was then employed to lower the impedance of gold contact sites to 100 kΩ(*42*).

To assemble rigid shuttle devices to assist NET implantation, pre-cut straight tungsten microwires (Advent Research Materials) at diameter of 50 µm were inserted into microconduits constructed of Polytetrafluoroethylene tubes (Sub-lite-wall tubing, O.D. 200 µm, I. D. 100 µm, Zeus) at a length of 4-6 mm. The microwires protruded the tube edge by 3–5 mm on both ends, while PEG solution was applied at the rear end of the microconduits to temporarily fix the microwires. The microconduit array was then mounted on the NET substrate using Epoxy (Loctite), and aligned under the stereomicroscope so that the lateral position of individual microwires best matched the positions of NET shanks and the microwires protruded the end of the NET shanks by 50-100 µm. The flexible segment of NETs was then attached onto the microwires using PEG solution as described elsewhere(*43*). The assembled device was ready for implantation after dried in air.

### Surgery and post-surgical preparation

Mice were anesthetized with isoflurane (3% for induction and 1% - 2% for maintenance) in medical O_2_. Body temperature was maintained at 37°C with a feedback-regulated heating pad (Far Infrared Warming Pad, Kent Scientific). Arterial oxygen saturation, heart rate, and breath rate were monitored via pulse oximetry (MouseSTAT, Kent Scientific). The animal was then placed supine in a stereotaxic frame (David Kopf Instrument). Carprofen (5 mg/kg) and dexamethasone (2 mg/kg) were administrated subcutaneously to reduce inflammation of the brain during the craniotomy and implantation procedure. The scalp was shaved and resected to expose skull between the bregma and lambda cranial coordinates. A circular portion of skull (about 3 mm in diameter) atop the somatosensory cortex was removed with a dental drill (Ideal Microdrill, 0.8 mm burr, Fine Science Tools) under constant sterile artificial cerebrospinal fluid (buffered pH 7.4) perfusion. Dura mater was partially removed to facilitate NET implantation. 4-shank, 32-contact NET 2D arrays at the inter-shank spacing of 250 µm were implanted stereotaxically using tungsten microwires as the shuttle device and bio-dissolvable adhesive as discussed in previous publication(*43*). After implantation of NET arrays at the vertical angle to the cortical surface, the carrier chip was carefully positioned and mounted on the remaining skull at 45° and about 5 mm away from the implantation sites to allow for optical access. A small hole was drilled on the contralateral hemisphere of the brain and a bare Ag wire was inserted into the brain as the grounding reference for electrical recording. A 3 mm round cover glass (#1, World Precision Instruments) was placed over the exposed, NET implanted brain area with a layer of artificial cerebrospinal fluid between the two. Gentle pressure was applied to the cover glass while the space between the coverslip and the remaining skull was filled with Kwik-sil adhesive (World Precision Instruments). An initial layer of C&B-Metabond (Parkell) was applied over the cyanoacrylate and the Kwik-sil. This process ensured a sterile, air-tight seal around the craniotomy and allowed for restoration of intracranial pressure. A second layer of Metabond was used to cement the coverslip and the NET carrier chip to the skull. A final layer of Metabond was used to cement a customized titanium head-plate for later head-constrained measurements.

Following surgery, animals were weighed daily for the first week and then once a week to ensure they did not fall below 90% of their free-feeding body weight. Additionally, analgesics (carprofen) and veterinary intervention were provided when some of the animals exhibited signs of pain and discomfort. The multimodal experiments started no sooner than four weeks after surgery to allow sufficient recovery of the animals. Animals were then handled and trained to head fixation in sessions of 20 min to an hour across the course of 3 - 5 days. Most animals were handled awake for habituation to head-fixation, while brief isoflurane anesthesia was used occasionally when the animal showed high level of anxiety during handling. No anesthesia was used for any animal from three days prior to baseline measurements to the conclusion of the experiments. Rose Bengal, a fast-clearing photothrombotic agent that photochemically triggers localized clot formation upon irradiation with green light(*44, 45*), was injected i.p. (50 μL, 30 mg/mL) in awake mice before the stroke induction session.

### Laser speckle contrast imaging and targeted photothrombosis

Laser speckle contrast imaging (LSCI) of CBF was performed using a 685 nm laser diode (50 mW, HL6750MG, Thorlabs) illuminating the craniotomy at an oblique angle to monitor the clot formation within the targeted area and the progression of ischemia. The backscattered laser light was relayed to a CMOS camera (acA1920-155μm, Basler AG) with 2 × magnification and acquired 1280 × 1024 pixel frames for a field of view (FOV) of 3.5 mm × 2.8 mm using custom software written in C++. The frame rate was 60 frames per second (fps) with a 5-ms exposure time. Simultaneously, laser light (200 mW, AixiZ) at 532 nm was patterned by a digital micromirror device (DMD, DLP3000, Texas Instruments). A small fraction of the light that scaled with the number of DMD pixels in the “on” state was delivered to the craniotomy to induce user-defined photothrombotic occlusions(*16, 46*) in the cortical vasculature using Rose Bengal. The projected DMD pattern was co-registered with the LSCI camera via an affine image transformation. This allowed for the selection of arbitrarily-shaped regions of interest using the LSCI imagery, which was then transformed into DMD coordinate space and loaded onto the device(*16*). Descending arterioles were the primary target because they serve as bottlenecks in the cortical oxygen supply(*47*). For all sessions of LSCI imaging and targeted photothrombosis, mice were awake and head-fixed on a customized, low-profile treadmill. Following photothrombosis, animals were closely monitored for weight loss and other signs of discomfort. Two animals out of twelve had post-stroke complications that were not successfully alleviated by veterinary interventions. They received humane euthanasia and were excluded from the study.

### Multi-exposure speckle imaging of Penumbral CBF

Another imaging system was used to perform multi-exposure speckle imaging (MESI) (*14, 15*) that longitudinally monitored CBF before and after stroke induction. A volume holographic grating wavelength-stabilized, single-frequency laser diode (785 nm, LD785-SEV300, Thorlabs) illuminated the craniotomy while simultaneously triggering 15 camera (acA1920-155 μm, 1920 × 1200 pixels, Basler AG) exposure durations ranging from 0.05 to 80 ms. The laser intensity was modulated such that the total amount of light was conserved across camera exposures to minimize the effects of shot noise. Backscattered light was collected by an objective (AF-S Nikkor 50 mm F/1.8G, Nikon) with a FOV of 2.92 mm × 4.02 mm and imaged onto the camera. CBF from the multiple pre-stroke sessions were average to provide the baseline values to normalize all the MESI imaging results from the same animal. For all MESI imaging sessions, mice were awake and head-fixed on a customized, low-profile treadmill.

### Electrophysiology

The LFP and extracellular spikes were recorded using 2D NET intracortical arrays at a sampling rate of 30 kHz with the bare Ag in the contralateral hemisphere of the brain as the grounding reference. The 2D NET array consists of 4 shanks, 4 × 8 contacts at inter-shank spacing of 250 µm, and contact center-to-center spacing of 80 µm on the shank. The implantation depth was controlled so that the shallowest contacts were about 100 µm below the pia layer, and the deepest contacts were about 660 µm deep. Voltage signals from the contacts were filtered between 0.5 Hz and 6000 Hz, amplified and digitized using a 32-channel RHD 2132 acquisition system (Intan Technologies). The impedance of all contacts was measured before each session, and contacts with impedance at 1 kHz larger than 2 MΩ were regarded as bad connections and excluded from the measurements. Electrical recordings were performed on awake animals consecutively with MESI before and after stroke induction session, and simultaneously during LSCI at the acute stroke induction session. For simultaneous LSCI and recording, the voltage pulse triggers the camera exposure in LSCI were recorded in the RHD 2132 acquisition system to synchronize both measurements. No photoelectric artifacts were detected during speckle imaging.

### Quantification and Statistical Analysis

#### Speckle imaging of CBF

For speckle imaging (LSCI and MESI), raw images captured by the camera were converted to speckle contrast images(*15, 48*) using a 7×7-pixel sliding window centered at every pixel of the raw image and an efficient processing algorithm allowing for real-time computation, display and data saving(*49*). During post-processing, speckle contrast images were averaged together (n = 45 for LSCI, n = 100 for MESI) and converted to inverse correlation time (1/τ_C_) images to provide a more quantitative measure of blood flow. For LSCI at the stroke induction session, each inverse correlation time image was then baselined against the first frame prior to blood flow reduction to calculate an estimate of relative change in blood flow (rCFB = τ_C,initial_/τ_C_)(*50, 51*). The same regions of interest defined for DMD structured illumination for photothrombosis were used to monitor the change of CBF during clot formation. For MESI, inverse correlation time images were aligned across multiple sessions through image registration, and 1/τ_C_ from the same regions of interest were repeatedly computed during the chronic time course(*14*). The values of inverse correlation time were then baselined against their average values from the pre-stroke sessions to calculate the relative change in blood flow over time (rCFB = τ_C,pre-stroke_avg_/τ_C_).

#### Quantification of LFP and spike sorting

For electrophysiology, a notch filter at 60 Hz was applied to the raw time traces at sampling rate of 30 kHz. The median signal from all channels was then used as the common mode reference and subtracted from each channel. Next, after high-pass filtering at 300 Hz, spike sorting was performed by thresholds detection and clustering in MountainSort(*41*), using similar settings and thresholds as reported previously. Adjacency radius was set to 100 to restrain the clustering neighborhoods to the immediate neighboring recording sites. The event detection threshold was set to 4.5 SD for all but one mouse (3.8 SD) that had multiple spikes of small amplitude (between 3.8 SD to 4.5 SD) appearing after stroke. Manual examination of mean waveforms of sorted units was then performed to reject noise clusters. To compute the total spike rate, the spiking events from all sorted units were grouped and combined by the contact on which the largest peak to peak amplitude of the units was detected. Spectrograms were performed on the LFP data using 1 s non-overlapping windows. Integrated power in different frequency bands, *i.e.*, 30 - 60 Hz and 60 – 110 Hz was calculated to extract the envelope of the signal. For acute sessions, LFP at all bands and spiking rate were normalized to the averaged value of the first 100 s prior to blood flow reduction. For longitudinal comparisons, a recording duration of 300 s was used to compute the mean and standard deviation as shown in Figure 4, and all values are normalized to the mean of all pre-stroke baseline sessions.

#### Quantification of neurovascular coupling coefficient

Time sequence of parenchyma CBF was determined from acute LCSI at the tissue where NETs were implanted at 1.33fps, and was interpolated at 1s intervals. Time sequence of neural data (LFP at a given band and spike rate) recorded by individual contacts were calculated using a 1-s time window. Neural data were then averaged among the selected groups of contacts on the same shank for a variety of depth profiles as described in the main text and supplementary figures. For each animal and each location studied, the time sequences of LFP, CBF and spike rate were sections to 1000 s sub-sessions. The cross-correlation between neural data and CBF was calculated using Matlab for the first 1000 s sub-session before the occlusion in the targeted arterioles formed. The temporal shift between the two sequences was constrained to [-1 5] seconds based on the knowledge of the temporal correlation between focal neural activation and the subsequence changes in CBF, and were found to be typically at 1 – 2 s (neural sequence was earlier than CBF responses). The same temporal shift was applied to offset the time sequences of CBF and neural data at later sub-sessions to compute the cross-correlation. Sub-sessions in which one or a few peri-infarct depolarizations (PIDs) occurred was excluded from the scatter diagram.

#### Tissue Processing and Analysis of Lesion Volume

Animals were overdosed with isoflurane and transcardially perfused with 0.1 M phosphate buffer saline (PBS) and 4% paraformaldehyde. Following perfusion, brains were frozen at −20°C for 30 – 60 min to allow separation from the skull without removing NETs from the brain. Brains were dissected and post-fixed *in situ* overnight in 4% paraformaldehyde at 4°C The brain tissue was then cryo-protected in 10% sucrose in paraformaldehyde until the tissue sunk (6 – 12 hrs), and then in 30% sucrose in paraformaldehyde for 48 hrs. The fixed tissue was then sliced into 30 µm thick coronal sections using a cryostat (Microm HM550). Every second section was mounted onto Gelatin subbed slides (SouthernBiotech), air dried overnight and Nissl stained with toluidine blue (Toluidine: Sodium Borate: dH_2_O in weight = 1:2:800) for lesion reconstruction and volume measurement(*17*). The lesion volume was measured as V = ΣA×T, where ΣA is the sum of the area of all the sections and T is the distance between section planes (90 μm)(*52*).

### Statistics

The CBF and neural data during the longitudinal tracking were statistically evaluated by two-way ANOVAs with Tukey’s post hoc test for multiple comparisons (OriginPro2015, OriginLab Corp.). NET contacts that were measured to have a high impedance (> 2 MΩ at 1 kHz) at any session during longitudinal tracking were identified as broken and were excluded from the statistics. This counted for 5% loss of the recording channels. Statistical test results are designated with n.s. when p-value > 0.05, one asterisk (*) when p-value < 0.05, two asterisks (**) when p-value < 0.01, three asterisks (***) when p-value < 0.001, and four asterisks (****) when p-value < 0.0001.

## Acknowledgments

We thank the Microelectronics Research Center at UT Austin for the microfabrication facility and support, the Animal Resources Center at UT Austin for animal housing and care, and CBRS Microscopy & Imaging Facility at UT Austin for histological processing.

## Author contributions

L.L., A.K.D. and C.X. conceived the experiments; F.H. fabricated the NET devices supervised by C.X. and L.L.; F.H. performed surgery and neural recording with helps from X.L. and Z.Z supervised by L. L.; F.H. and C.S. performed *in vivo* imaging experiments supervised by L.L. and A.K.D.; F.H. and M.R.W. performed histology supervised by L.L. and T.A.J.; F.H., H.Z., and C.S. analyzed data supervised by L.L., A.K.D. and T.A.J.; L.L and F.H. wrote the manuscript with input and revisions from all authors.

## Data and materials availability

All data needed to evaluate the conclusions in the paper are present in the paper and/or the Supplementary Materials. Additional data available from authors upon request.

