## Supplementary information for "Multimodal mapping of neural activity and cerebral blood flow reveals long-lasting neurovascular dissociations after small-scale strokes"

Fig. S1. Representative recordings from a 4-shank, 32-contact NET implant in an awake head-fixed mouse.

Fig. S2. Formation of an infarct as monitored by continuous LSCI of CBF.

Fig. S3. LFPs at multiple frequency bands consistently disassociate with CBF under ischemia.

Fig. S4. The neurovascular disassociation with ischemia is consistent within a variety of depth profiles of neural activity.

Fig. S5. Neuronal hyperexcitability is sparse and occurs days after hyper-perfusion in the surrounding tissue.

Fig. S6. Longitudinal neurovascular disassociation at additional LFP frequency bands.

Fig. S7. Lack of correlation between the severity of neurovascular disassociation and the area of severe CBF deficits (A), and between final CBF and acute CBF or LFP (B, C).

Table S1. Experimental subject numbers.

Movie S1. LFP at 60 – 110 Hz and spike firing rate (FR) from all NET contacts simultaneously recorded with full-field LCSI of CBF at the acute stroke session.

Movie S2. Longitudinal MESI of CBF from pre-stroke baselines to Day 55 after stroke, and measurements of LFP at 60 –110 Hz and spike rate of all NET contacts at the same days.

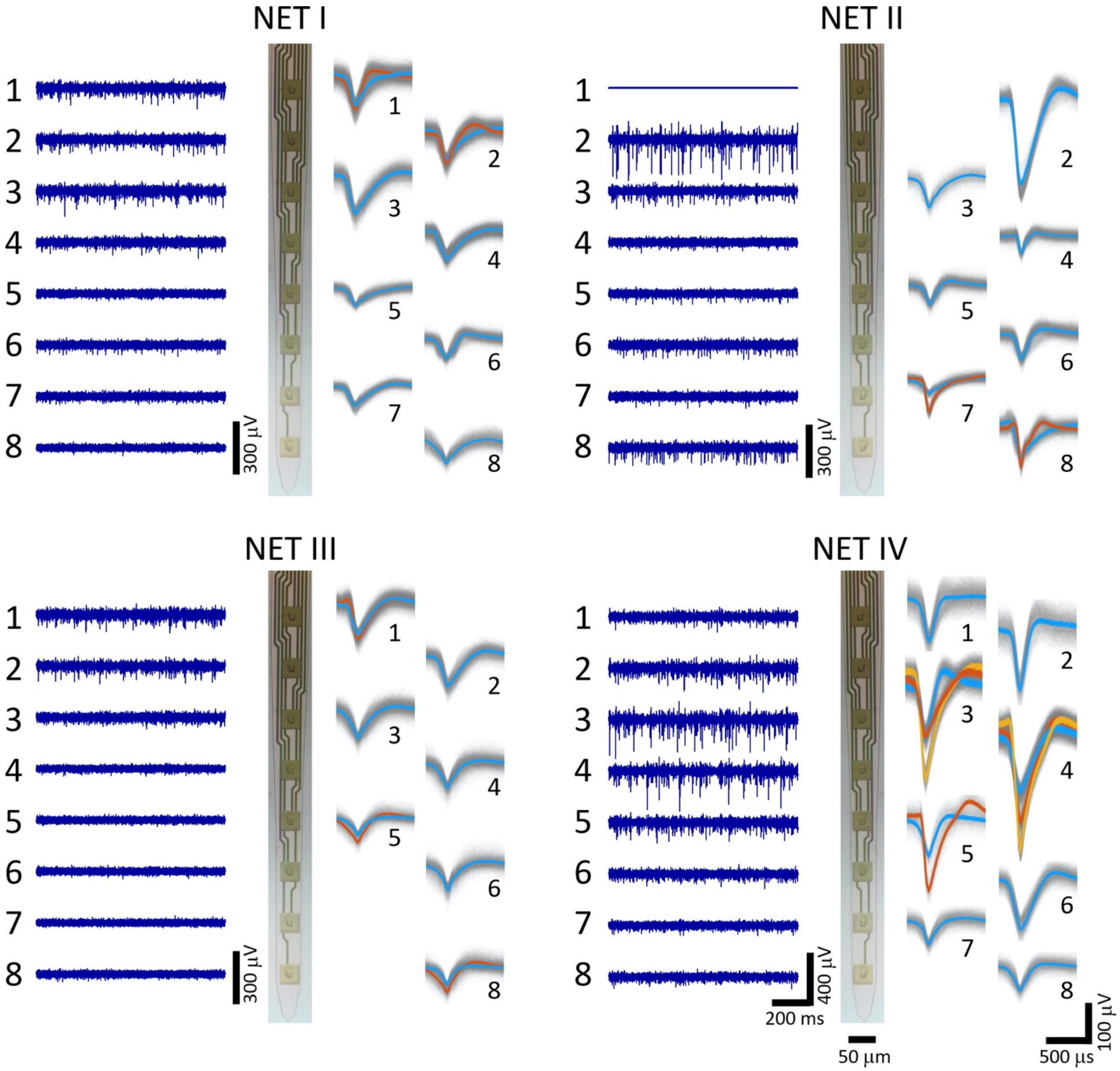


**Figure S1. Representative recordings from a 4-shank, 32-contact NET implant in an awake head-fixed mouse.** Each panel shows representative recording traces high-pass filtered at 300 Hz (left), photo of the NET shank (middle), and sorted spike waveforms by individual contacts (right).

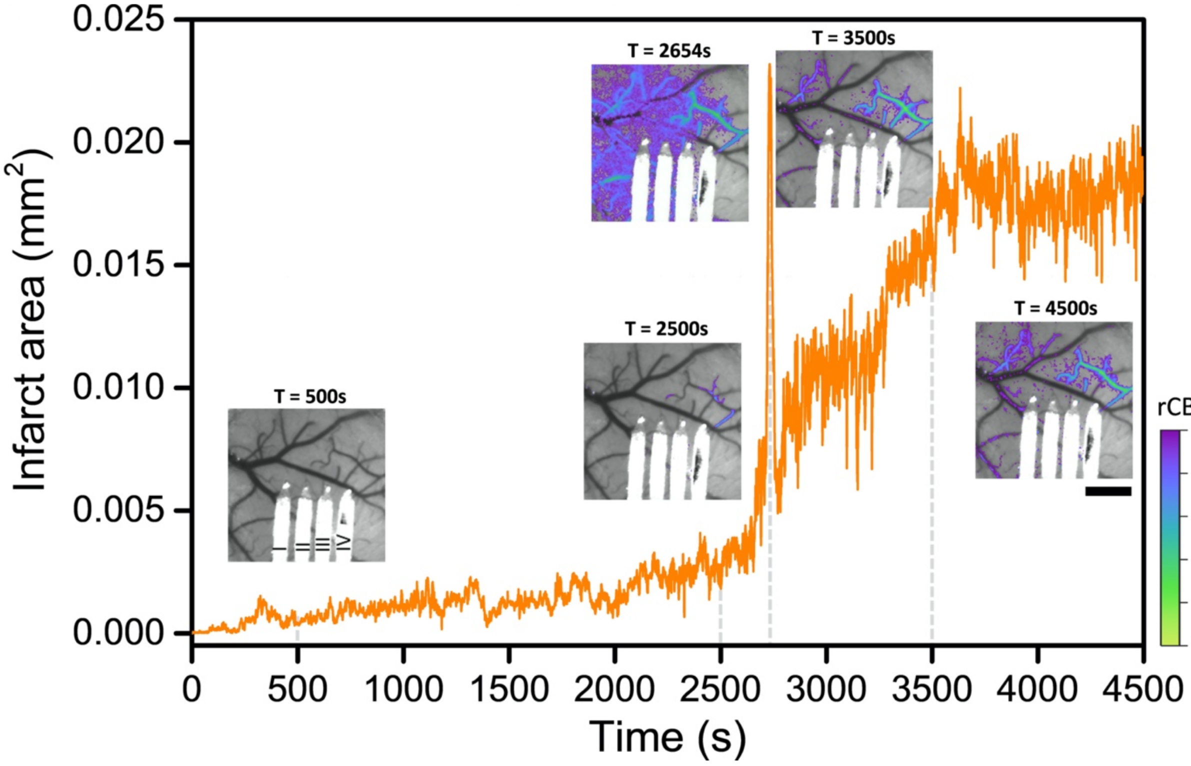


**Figure S2. Formation of an infarct as monitored by continuous LSCI of CBF.** The main panel plots the time-dependent infarct size determined as the area with relative CBF < 20% of the pre-stroke baseline. The sharp peak at 2654 s marks the occurrence of a peri-infarct depolarization (PID). Insets show the representative time series of LSCI of CBF in the same animal. Color map overlays the relative CBF < 50% baselined against pre-photothrombotic images. Scale bar: 500 μm.

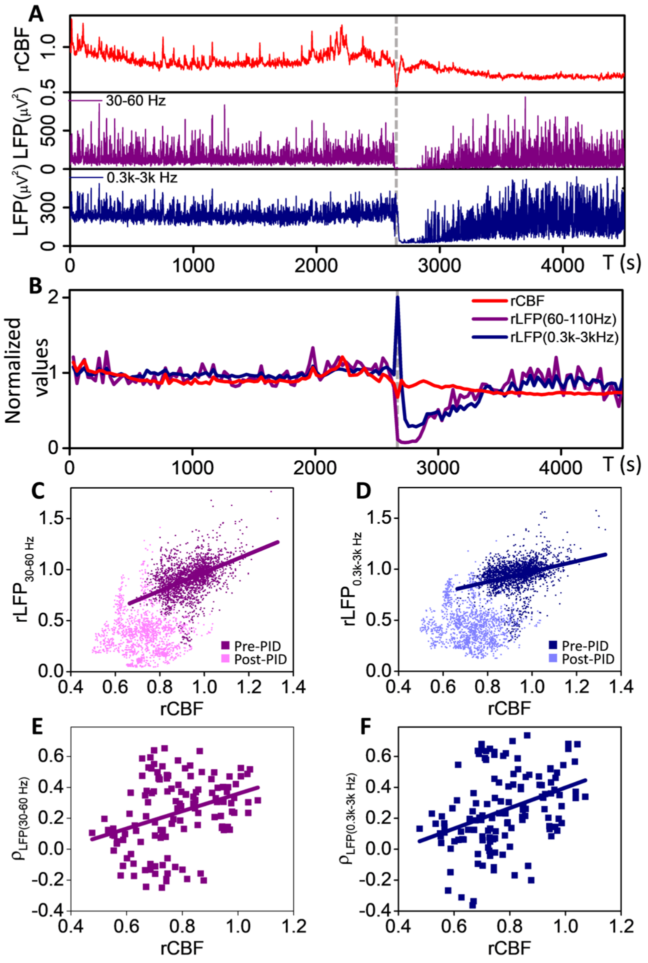


**Figure S3. LFPs at multiple frequency bands consistently disassociate with CBF under ischemia. (A, B)** Simultaneous measurement of CBF and LFP at 30-60 Hz and 0.3k - 3k Hz in the same acute stroke session and using the same contact (NET IV-4) as in **Fig. 2E.** Normalized and 30s- averaged values were used in **(B)** to visualize the neurovascular dissociation. Dashed lines mark a PID event that was excluded from the following analysis. **(C, D)** Scatter diagrams of rCBF and rLFP at 30-60 Hz **(C)**, and at 0.3k-3k Hz **(D)**. Similar to **Fig. 2**, the positive linear correlation (darker color) breaks down as ischemia developed (lighter color) after the spontaneous occurrence of PIDs. Pearson's correlation was applied. $\rho = 0.43$, p-value = 5.5e-98 **(C)**; $\rho = 0.30$, p-value = 6.3e-47 **(D)**. **(E, F)** Correlation coefficient computed between rCBF and rLFP at 30-60 Hz (**E**) and at 0.3k-3kHz (**F**) for all locations and animals. Pearson's correlation was applied. $\rho = 0.34$, p-value = 8.4e-5 **(E)**; $\rho = 0.37$, p-value = 1.1e-5 **(F)**.

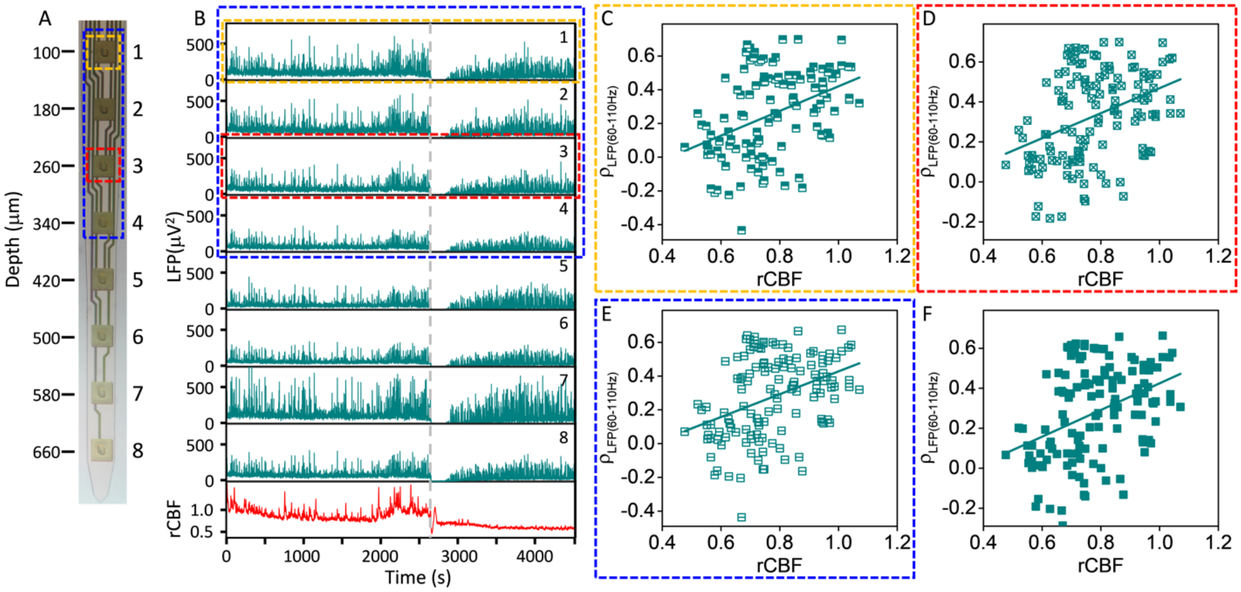


**Figure S4. The neurovascular disassociation with ischemia is consistent within a variety of depth profiles of neural activity. (A)** Photo of a NET shank marking the estimated implantation depth of all contacts. Dashed boxes enclose contacts used to compute neural activity in **(C-F)**. **(B)** LFPs from individual contacts and the relative CBF from the nearby tissue. **(C-F)** The coupling coefficients computed between CBF and LFP using **(C)** the shallowest contact; **(D)** the contact that had the largest correlation coefficient at pre-stroke; **(E)** the averaged value of a subset of the shallower contacts (Contacts 1-4); and **(F)** the entire shank as used in the main text. Pearson's correlation was applied to determine the dependence of the coupling coefficient on CBF reduction. $\rho$ = 0.40, p-value = 4.1e-6 **(C)**; $\rho$ = 0.38, p-value = 8.5e-6 **(D)**; $\rho$ = 0.39, p-value = 4.9e-6 **(E)**; $\rho$ = 0.40, p-value = 3e-6 **(F)**.

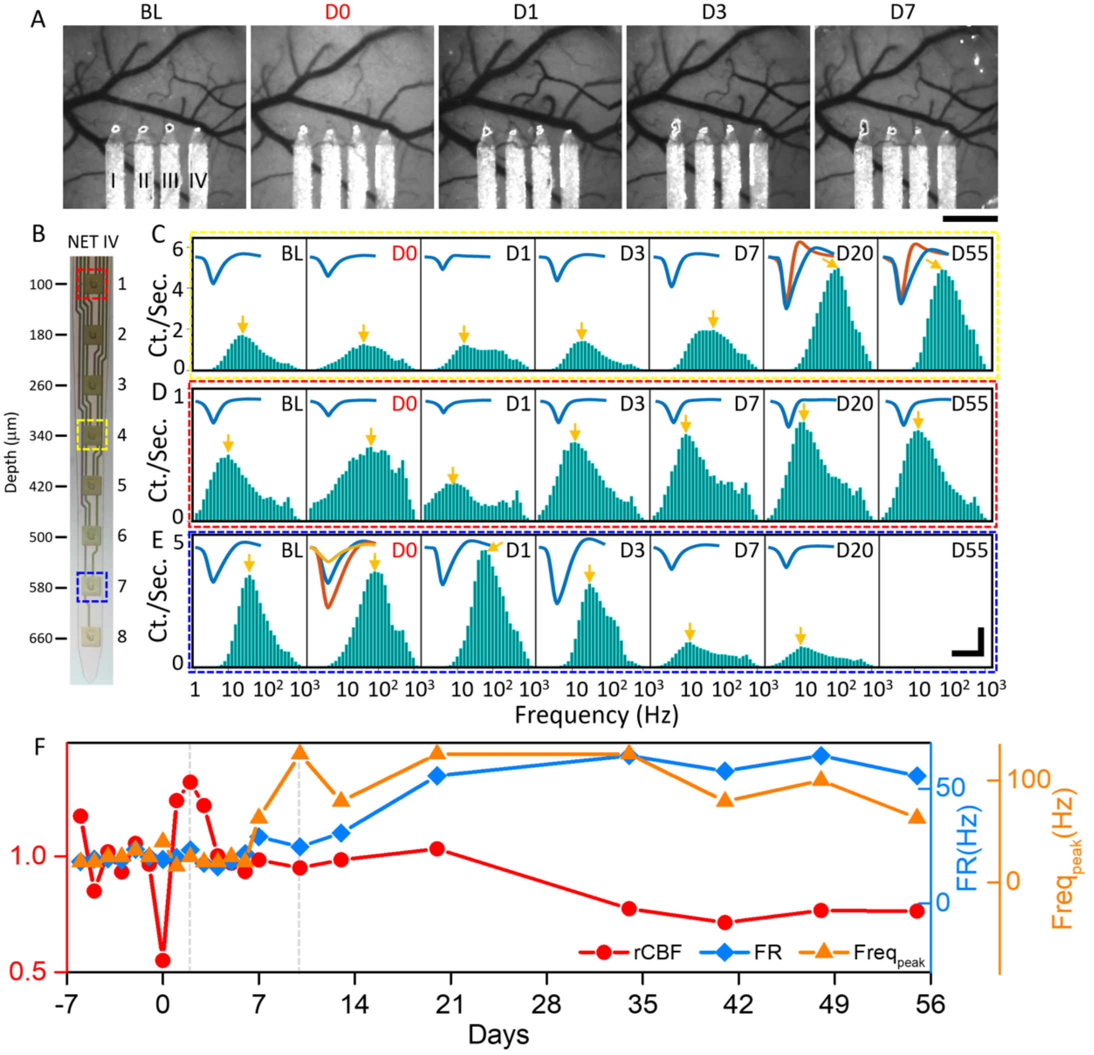


**Figure S5. Neuronal hyperexcitability is sparse and occurs days after hyper-perfusion in the surrounding tissue. (A)** MESI of CBF at multiple temporal coordinates before and after stroke showing hyper-perfusion happened at Day 1 after stroke induction. NET shanks are marked in the pre-stroke baseline (BL) image. D0: immediately after stroke**. (B)** Photo of a NET shank. Color-coded dashed boxes mark the contacts used to record neural activity in **(C-E)**. **(C-E)** Distribution of discharge frequencies in unit recordings at multiple temporal coordinates before and after stroke from the contacts marked in **(B)**, showing the distinct patterns of excitability with time: hyperexcitability, contact 4 **(C)**; little change in excitability, contact 1 **(D)**; and hypoexcitability, contact 7 **(E)**. Insets show the spike waveforms. Colors denote the putative units. Arrows mark the peak frequency in the distribution. **(F)** Time dependence of the averaged firing rate (blue diamond)

and the peak frequency in the frequency distribution of unit firing rate (orange triangle) recorded by contact 4 in **(C)** overlaid with relative CBF (red circle). Neuronal hyperexcitability occurs at a later time than hyper-perfusion. Scale bars: 500 μm **(A)**, 100 μV (vertical in **C-E**) and 1 ms (horizontal in **C-E**).

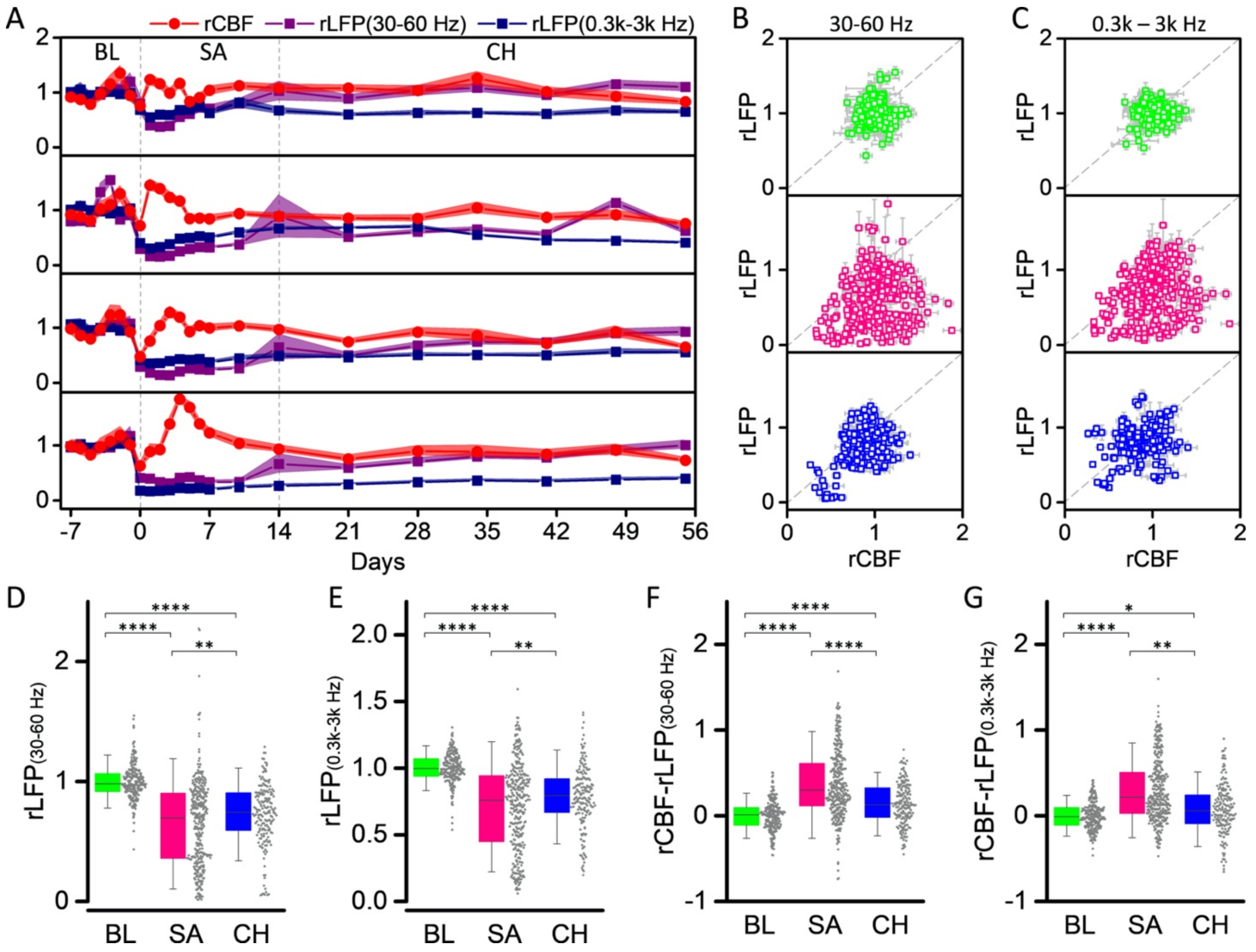


**Figure S6. Longitudinal neurovascular disassociation at additional LFP frequency bands.** Same animals as in **Fig. 3** in the main text. **(A)** Relative CBF, LFPs at 30-60 Hz and 0.3k-3k Hz at the 4 NET locations baselined against pre-stroke values. Solid dots present the mean value, and shading shows the STD. **(B, C)** Scatter diagrams between LFP (30-60 Hz) and CBF **(B)**, and between LFP (0.3k–3k Hz) and CBF **(C)** in three phases: baseline (BL), consecutive 7-days daily measurements before stroke; subacute (SA), Day 0 – 14 post stroke; and chronic (CH), week 3 – 8 post-stroke. All values are normalized to the mean of the baseline values. **(D-G)** Box plots showing the relative values of LFP (30-60 Hz) **(D)**, LFP (0.3k-3k Hz) **(E)**, and their difference CBF-LFP (30-60 Hz) **(F)** and CBF- LFP (0.3k-3k Hz) **(G)** at the three phases: baseline (green); sub-acute (magenta); chronic (blue). All data plotted as dots. Significant levels shown as: *, p-value < 0.05; **, p-value < 0.01; ***, p-value < 0.001; ****, p-value < 0.0001.

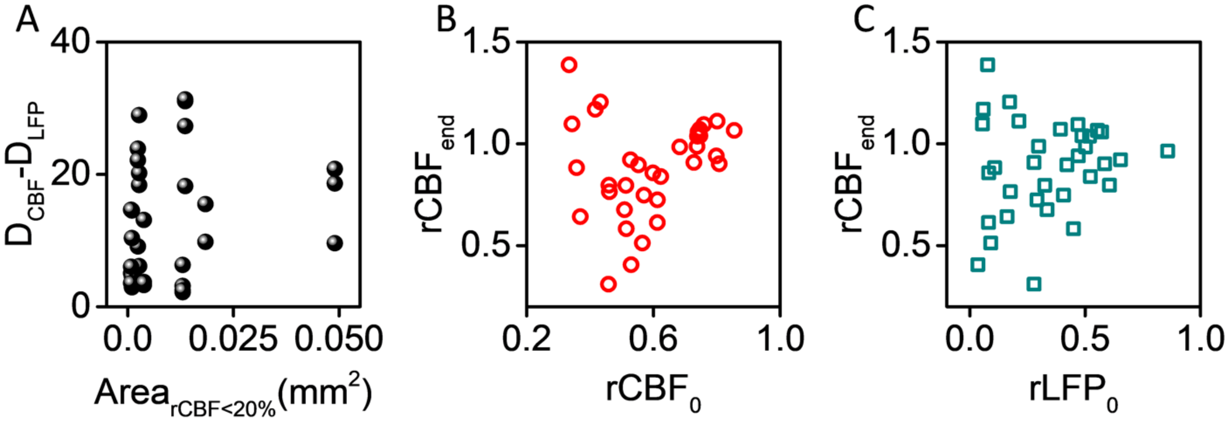


**Figure S7. Lack of correlation between the severity of neurovascular disassociation and the area of severe CBF deficits (A), and between final CBF and acute CBF or LFP (B, C).** Pearson's correlation was used. $\rho = 0.21$, *p*-value = 0.25 in **(A)**; $\rho = 0.20$, *p*-value = 0.25 in **(B)**; $\rho = 0.15$; *p*-value = 0.41 in **(C)**.

**Table S1. Experimental subject numbers.**

| | | Animal | Location | Animal | Location | |
|---|---|---|---|---|---|---|
| **Total** | | **12** | **48** | | | |
| **Experiment** | | **In use** | | **Exclusion** | | **Reason of exclusion** |
| Surgery | | 12 | 47 | 0 | 1 | 1 NET shank failed to be implanted intracortically |
| Acute study | | 10 | 39 | 2 | 8 | High level of locomotion induced artifacts |
| Chronic study | Fig. 4 | 10 | 37 | 2 | 8 | Animal loss due to stroke complication |
| | | | | 0 | 2 | Tissue-sinking damaged NET shank |
| | Fig. 6 | 9 | 33 | 1 | 4 | Window clouding at week 5 |

**Movie S1.** LFP at 60 – 110 Hz and spike firing rate (FR) from all NET contacts simultaneously recorded with full-field LCSI of CBF at the acute stroke session. The occurrence and propagation of a peri-infarct depolarization event led to the silence of neural activity and the expansion of the ischemic area.

**Movie S2.** Longitudinal MESI of CBF from pre-stroke baselines to Day 55 after stroke, and measurements of LFP at 60 –110 Hz and spike rate of all NET contacts at the same days. Same Mouse as in **Fig. 3**.
